# Detection of Mid-parent Heterosis Genes in Large-Scale Unreplicated RNA-Seq Experiments

**DOI:** 10.64898/2025.12.11.693571

**Authors:** Yunhui Qi, Peng Liu

**Affiliations:** Department of Data Science, Dana-Farber Cancer Institute; Department of Statistics, Iowa State University

**Keywords:** Dispersion estimation, Gene expression heterosis, Mid-parent heterosis, Negative binomial distribution

## Abstract

Mid-parent heterosis (MPH), characterized by hybrid trait values deviating from the mid-parent average, is a well-documented phenomenon whose genetic basis remains poorly understood. Identifying genes associated with MPH is crucial for uncovering the molecular mechanisms underlying heterosis. Recent large-scale RNA-sequencing (RNA-seq) experiments enable the evaluation of heterosis genes across numerous families; however, replication is often infeasible due to cost and labor constraints, resulting in unreplicated large-scale datasets and posing statistical challenges for dispersion estimation and reliable inference. To address this issue, we propose a novel two-stage likelihood ratio test (2sLRT) for detecting MPH genes in unreplicated RNA-seq experiments. In the first stage, genes and families without evidence of differential expression across varieties are identified, and the corresponding varieties are used as pseudo-replicates to estimate dispersion. In the second stage, a likelihood ratio test based on the negative binomial distribution is employed to test for MPH. Simulation studies demonstrate that 2sLRT achieves higher power and better false discovery rate control compared to existing approaches. Application of 2sLRT to a maize RNA-seq dataset with 599 families further highlights the method’s effectiveness in revealing meaningful patterns of MPH gene expression.

## 1 Introduction

Heterosis, or hybrid vigor, is a striking biological phenomenon in which hybrid offspring outperform their inbred parents, displaying enhanced growth, yield, or resilience. Widely documented in both plant (Wakchaure et al., 2015) and animal breeding (Stuber and Janick, 2010), heterosis has long fueled progress in agriculture and breeding. Yet, despite its enormous practical value, the genetic basis of heterosis remains complex and only partially understood (Wu et al., 2021). With the advent of high-throughput technologies such as microarrays and RNA sequencing (RNA-seq), researchers have begun to uncover molecular footprints of heterosis at the gene expression level—a phenomenon referred to as gene expression heterosis. Importantly, gene expression heterosis has been proposed as one of the mechanisms contributing to phenotypic heterosis (Swanson-Wagner et al., 2006; Springer and Stupar, (2007). Depending on how hybrid expression compares to parental levels, gene expression heterosis is typically categorized into three forms: mid-parent heterosis (MPH), where expression in the hybrid deviates from the parental mean; high-parent heterosis (HPH), where the hybrid exceeds both parents; and low-parent heterosis (LPH), where the hybrid falls below both parents.

Several Bayesian methods have been developed to identify heterosis genes using microarray and RNA-seq data. Ji et al. (2014) proposed an empirical Bayes approach based on a mixture distribution to detect heterosis genes with microarray data. For RNA-seq count data, Niemi et al. (2015) formulated a hierarchical model based on the negative binomial distribution to identify HPH and LPH genes, also through empirical Bayes methods. While empirical Bayes analysis offers computational efficiency compared to fully Bayesian analysis, its success relies on the accurate estimation of hyperparameters. Landau et al. (2018) proposed a general hierarchical model for fully Bayesian analysis that eliminates the need for hyperparameter estimation. However, all these Bayesian methods impose parametric assumptions on the hyperparameters, which may not fit data well. To address this limitation, Bi and Liu (2023) proposed a semi-parametric empirical Bayes approach by directly modeling the fold changes between each parent and the hybrid with a Dirichlet process. This semi-parametric approach offers flexibility and potentially more accurate modeling of heterosis genes, but is very computationally intensive.

Recently, some large-scale, multi-family RNA-sequencing experiments have been carried out to simultaneously evaluate heterosis using numerous families, with each family containing three varieties: two parental varieties and one hybrid. While such experiments enable the evaluation of heterosis across diverse genetic backgrounds, it is cost- and labor-prohibitive to include replicates, resulting in unreplicated, large-scale RNA-seq experiments (Xu et al., 2025). However, all the aforementioned methods were proposed to analyze datasets from a single family with several biological replicates for each variety. Although Poisson models do not require replicates to fit the model parameters and draw inference on heterosis genes, Poisson distributions do not handle over-dispersion, a common characteristic observed in RNA-seq data, and will likely introduce a lot of false positives. The negative binomial distribution has been a standard modeling approach for RNA-seq count data due to its ability to handle over-dispersion. However, negative binomial models require replicates to estimate their model parameters: mean and dispersion. Currently, there are no available statistical methods that analyze RNA-seq data from unreplicated multi-family RNA-seq heterosis experiments and handle overdispersion. To fill this gap, we propose a two-stage likelihood ratio test (2sLRT) based on negative binomial models to identify MPH genes within each of all families.

The core structure of our proposed 2sLRT method is described in Figure 1. In the first stage, we identify some “null families” whose varieties do not exhibit differential expression, and use null families as pseudo-treatments and varieties in each null family as pseudo-replicates to estimate dispersion parameters for each gene. In the second stage, we perform a negative binomial likelihood ratio test (LRT) to assess MPH for each gene-family combination, conditional on the estimated dispersion parameters. Our approach does not require prior assumptions as in Bayesian methods, is computationally efficient, and enables robust testing for MPH genes in large-scale, unreplicated RNA-seq experiments. Simulation studies demonstrate that our method effectively controls the false discovery rate (FDR) and has higher power than Poisson-based LRT for MPH detection and a naive estimation method in practice. When applied to a maize experiment with 599 families, our method controls false positives and identifies biologically meaningful heterosis genes.

**Figure 1.**
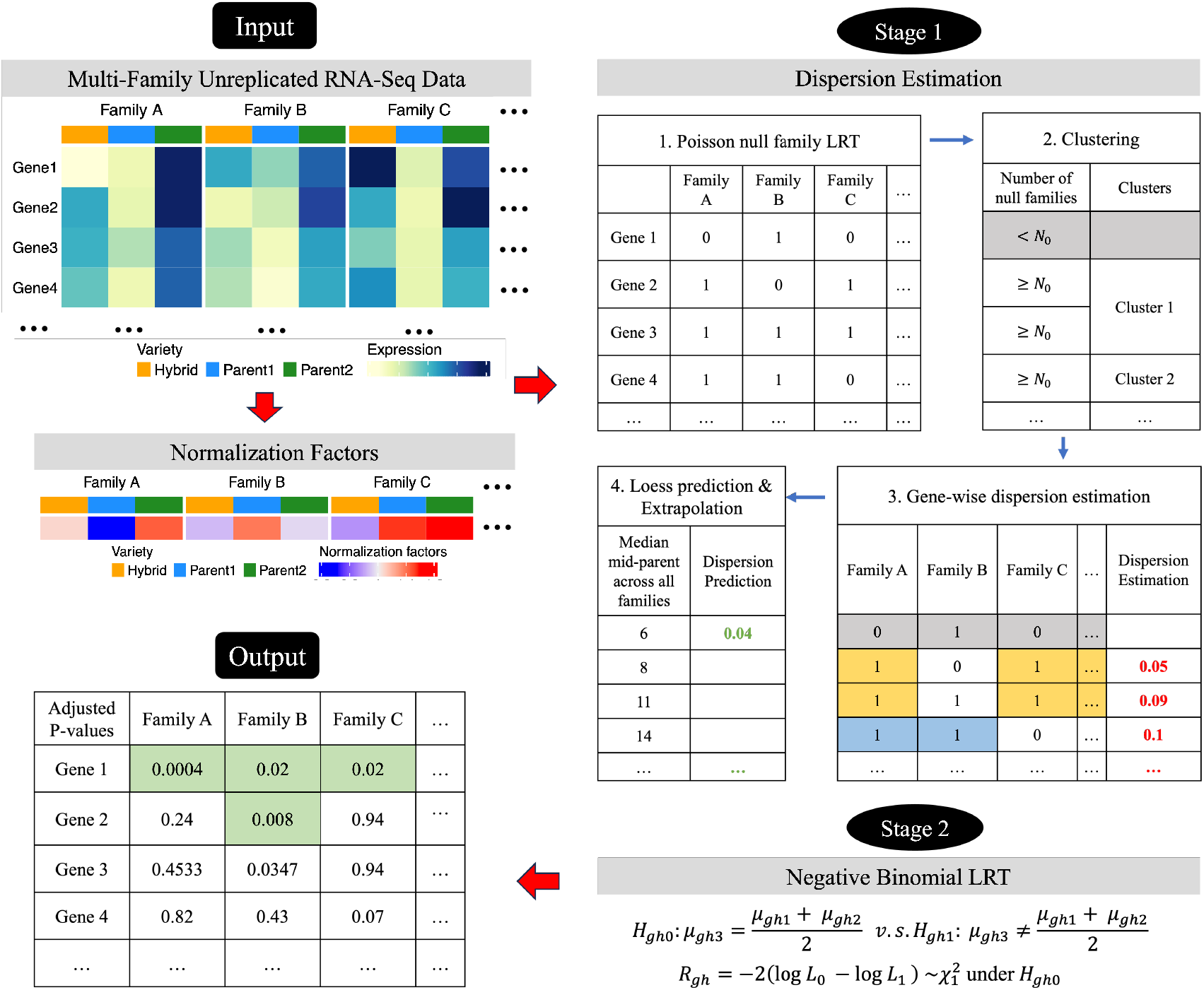
Schematic framework of the two-stage likelihood ratio test (2sLRT) for identifying mid-parent heterosis (MPH) genes in multi-family, unreplicated RNA-seq datasets (example dataset shown for illustration).

The remainder of the paper is structured as follows. In Section 2, we introduce the methodology and computational algorithms underlying the proposed 2sLRT. Section 3 reports simulation studies to assess the performance of 2sLRT. In Section 4, we apply our method to a multi-family, unreplicated maize gene expression dataset. Finally, Section 5 provides some concluding remarks and discussion. An R package for implementing 2sLRT can be accessed from the GitHub repository https://github.com/yunhuiqistat/TwoStageLRT.

## 2 Methods

In this section, we begin by assuming known dispersion parameters and constructing a likelihood ratio test based on negative binomial models to determine whether a gene exhibits MPH within a single family. Next, we outline a strategy for estimating dispersion parameters by borrowing information across multiple families and genes. Finally, we present the complete algorithm for detecting MPH genes in large-scale, unreplicated RNA-seq experiments.

### 2.1 Negative binomial LRT with known dispersion

In an unreplicated RNA-seq experiment, let *y*_*ghi*_ denote the observed count for gene *g* and variety *i* in family *h*, where *i* = 1, 2, 3 corresponds to the three varieties: parent one, parent two, and the hybrid offspring; and *g* = 1, …, *G* and *h* = 1, …, *H*, where *G* and *H* represent the total numbers of genes and families, respectively. We assume that *y*_*ghi*_ follows a negative binomial distribution:

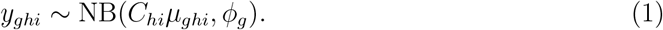

*C*_*hi*_ is the normalization factor for the sample corresponding to variety *i* in family *h*, and *C*_*hi*_ adjusts technical differences between samples, such as sequencing depth. Several R packages, such as edgeR (Robinson et al., 2010) and DESeq (Anders and Huber, 2010), offer robust methods to estimate the normalization factors. Once estimated, these normalization factors are treated as known in the subsequent analysis, a common practice in statistical analyses involving RNA-seq data. The parameter *µ*_*ghi*_ is the normalized mean expression level for gene *g*, family *h* and variety *i*; and *φ*_*g*_ is the dispersion parameter for gene *g*. Note that we assume the same dispersion parameter for each gene across varieties, consistent with the gene-specific dispersion assumption commonly used in differential expression analysis (Anders and Huber, 2010; Robinson et al., 2010).

Under model (1), gene *g* is said to be MPH in family *h* if the normalized mean expression level of the hybrid *µ*_*gh*3_ is not equal to the average of the normalized mean expression levels across the two parents. Accordingly, to test whether gene *g* is MPH or not in family *h*, we perform a hypothesis test for :

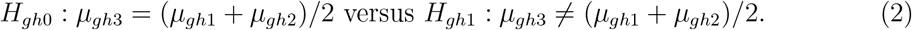

Assuming normalization factors *C*_*hi*_ and dispersion parameter *φ*_*g*_ are known, we have the following log likelihood of *µ*_*ghi*_:

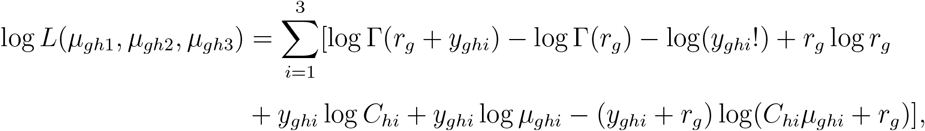

where *r*_*g*_ = 1/*φ*_*g*_ and Γ(·) denotes the Gamma function. The maximum likelihood estimator (MLE) of *µ*_*ghi*_ can be easily derived under the alternative hypothesis as 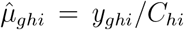 for each variety. The MLE of *µ*_*ghi*_ under *H*_*gh*0_, denoted by 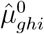, does not have a closed form. Numerical algorithms, such as the Broyden method, the full Newton method, or the Newton-Raphson method, are required to solve the system of equations:

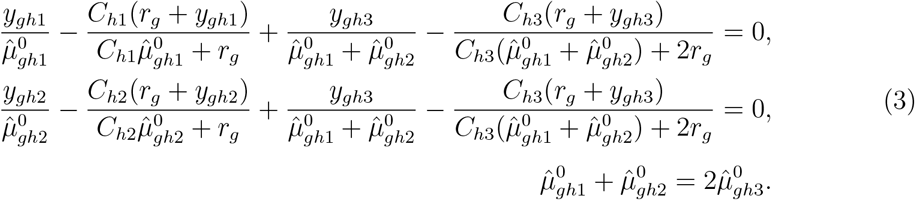

With the MLEs, gene *g* will be declared to be MPH in family *h* if the following likelihood ratio test statistic

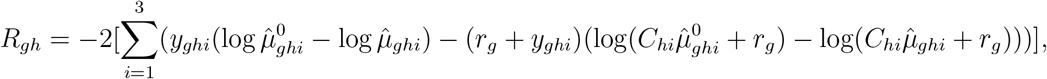

is bigger than 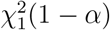 at the significance level *α*.

### 2.2 Estimation of dispersion parameters

In practice, the dispersion parameter for each gene is unknown and must be estimated from the data. Existing methods require biological replicates at the variety level, making them unsuitable for the unreplicated designs. To overcome this limitation, we propose a multi-step strategy for estimating dispersion parameters. The central idea is to identify genes and families that are not differentially expressed across varieties, so that varieties within these null families can be treated as pseudo-replicates. This enables the application of established methods that borrow information across genes to obtain stable dispersion estimates.

The first step of our procedure is to perform a Poisson likelihood ratio test (LRT) to identify, for each gene, families that show no evidence of differential expression across varieties, which we term “null-families” for the corresponding genes. The Poisson model, characterized by a single rate parameter, allows for maximum likelihood estimation under both the null and alternative hypotheses—even in the absence of biological replicates—making it well-suited for unreplicated RNA-seq data. The resulting *p*-values are adjusted for false discovery rate (FDR) control using the Benjamini-Hochberg method (Benjamini and Hochberg, 1995), producing a matrix 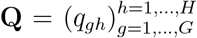, where *q*_*gh*_ is the adjusted *p*-value for gene *g* in family *h*. We acknowledge that such a Poisson-based method on over-dispersed data tends to produce false positives (Anders and Huber, 2010). Therefore, this step can be regarded as a conservative method to select the null families.

For each gene, the null families identified by Poisson LRT can be viewed as pseudo-treatments within which the three non-differentially expressed varieties are considered as pseudo-replicates. Pooling all null families for each gene, we could estimate *φ*_*g*_, the dispersion parameter for each gene. Such per-gene estimates do not utilize information from other genes. However, methods borrowing information across genes have shown effective improvement on dispersion parameter estimation (Anders and Huber, 2010; Robinson and Smyth, 2008). To borrow information across genes, we identify genes that share the same set of null families and apply existing methods such as edgeR and DESeq2. Ideally, we can get a partition of genes so that each partition gives a dataset with a sufficient number of genes to borrow information across genes, and a sufficient number of common null families to ensure a large enough sample size. This turns out to be a challenging task because (i) many genes share few, if any, similar null families with other genes, limiting opportunities for grouping based on shared structure; and (ii) some null families are associated with only a small number of genes, reducing the potential benefits of borrowing information across genes. The accuracy of dispersion estimation depends on the sub-matrix size, specifically, the number of selected genes and the number of null families. As a practical way to tackle these challenges, we introduce a clustering-based method for selecting subsets of genes and families.

The second step of our procedure, clustering, begins with an indicator matrix **I** denoting the null families for each gene. In this matrix, an entry in the *g*-th row and *h*-th column is 1 if the *h*-th family is declared a null family for gene *g*, and 0 otherwise, after controlling FDR at a user-specified level *α*_0_. Since genes with only a few null families provide limited sample sizes, we retain only those genes whose number of null families exceeds a threshold *N*_0_ in cluster analysis, thereby ensuring sufficient sample size in the subsequent estimation step. To group genes with similar patterns of null family distribution, we calculate similarity across genes based on the Jaccard distance (Jaccard, 1912) from the indicator matrix, and perform hierarchical clustering with complete linkage to group the genes into *K* clusters.

The third step estimates dispersion parameters for genes in each cluster. For each cluster, we identify the gene with the lowest count of null families, which typically corresponds to the most shared null families, and use the corresponding null families as the pseudo-treatment groups to estimate dispersion parameters for genes in the cluster. For such “pseudo-replicated” data, there are several well-performing methods available that borrow information across genes, such as edgeR.

In the final step, we estimate dispersion parameters for genes excluded earlier (genes with a null family count below the threshold *N*_0_). To do this, we first fit a loess curve to the dispersion estimates and the median of mid-parent expression levels across families using genes included in the previous estimation steps. Here, the mid-parent expression level is defined as the average of the expression levels across the two parental varieties. For the remaining genes, the dispersion parameter is estimated by evaluating this curve at their median mid-parent expression levels, a strategy similar to the ones used in DESeq and edgeR (Anders and Huber, 2010; Robinson et al., 2010). If a gene has a median mid-parent expression level beyond the fitted range, the estimate from the nearest median mid-parent expression level is used to extrapolate its dispersion.

A detailed step-by-step implementation of our proposed method for dispersion parameter estimation is described in Algorithm 1 of the Supplementary Materials.

### 2.3 Two stage LRT

Our proposed 2sLRT algorithm (Figure 1) combines the stage of dispersion parameter estimation and the stage of LRT based on negative binomial models to detect MPH genes with unreplicated RNA-seq experiments.

Given an RNA-seq data with *G* genes and *H* families—each with three varieties—we first estimate normalization factors across samples. Next, our 2sLRT procedure begins by applying the Poisson LRT to identify null families for each gene. Dispersion parameters are then estimated using Algorithm 1 in the Supplementary Materials. With these estimates, a negative binomial LRT is performed to detect MPH, while controlling FDR via the Benjamini-Hochberg procedure. Genes exhibiting MPH in each family are identified based on the specified FDR threshold. The full procedure is detailed in Algorithm 2 in the Supplementary Materials.

## 3 Simulations

In this section, we evaluate the performance of our proposed 2sLRT using simulation studies. We simulated RNA-seq data using a hierarchical structure, with the workflow visually outlined in Figure S1(a) of the Supplementary Materials. More specifically, for each gene-family combination, the means for parental lines (*µ*_*gh*1_ and *µ*_*gh*2_) are simulated from a gamma distribution with a gene-specific rate parameter simulated from a uniform distribution. Given the parental means, the hybrid mean is set to be 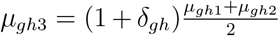, where *δ*_*gh*_ defines the effect size of MPH for gene *g* in family *h*, and *δ*_*gh*_ = 0 corresponds to genes without MPH. The normalization factors *C*_*hi*_ are simulated from a uniform distribution with the support of 0.8 to 1.3. The dispersion parameter *φ*_*g*_ is simulated according to a function of the mid-parent mean (*mp*) plus a random effect *E*_*g*_ from a normal distribution, where 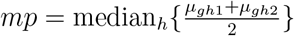. The function of *mp* is chosen based on the fitted relationship between estimated *φ*_*g*_ and *mp* from the dataset to be described in Section 4. In rare cases when the simulated *φ*_*g*_ is negative, it is replaced with 0. With the simulated mean expression levels (*µ*_*ghi*_), heterosis effect sizes (*δ*_*gh*_), normalization factors (*C*_*hi*_) and dispersion parameters (*φ*_*g*_), the negative binomial counts are simulated according to model (1).

The effect size of MPH, *δ*_*gh*_, can be positive, negative, and or zero (non-heterosis). Two signal strengths are considered: a strong signal with *δ*_*gh*_ = 3 or −3/4, where the hybrid mean is four times or one-quarter of the mid-parent mean, and a weak signal with *δ*_*gh*_ = 2 or −2/3, corresponding to hybrid mean being three times or one-third of the mid-parent mean.

Analysis of real data shows that some genes exhibit MPH in more families than others. To reflect this heterogeneity, our simulations consider two patterns of MPH occurrence based on the proportion of MPH families per gene. In pattern A, each gene has a fixed proportion of MPH families, though the specific families differ across genes: one group of genes shows MPH in 90% of families, and another in 5%, representing high- and low-MPH contrasts.

In pattern B, the proportion of MPH families varies by gene, drawn from the empirical distribution of gene-wise proportions estimated from real data. Details are illustrated in Figure S1(b) of the Supplementary Materials.

In total, we have 8 simulation settings combining different dimensions (*G* = 10000, *H* = 150; and *G* = 30000, *H* = 600), different heterosis gene patterns (A and B), and different heterosis effect sizes (strong and weak). For each setting, we simulated 50 datasets and applied the proposed 2sLRT to each of them and compared its performance with two other methods. One method is a current practice by biologists corresponding to a point estimate of MPH ratio, i.e., the absolute difference between the hybrid mean and the mid-parent mean divided by the mid-parent mean. We will refer to this method as the point estimation method. The other method is a Poisson LRT testing the hypothesis (2) to detect MPH genes. Derivation of the Poisson LRT for MPH detection is given in Section S2 of Supplementary Materials.

In the main text, we present simulation results for weak signals with *G* = 10000, *H* = 150 and equal parental means (*µ*_*gh*1_ = *µ*_*gh*2_) in Table 1 and Figure 2. Additional results for equal parental means – including weak signals with *G* = 30000, *H* = 600, and strong signals for both dimension settings are presented in Section S3.2 of Supplementary Materials. Results for all eight settings under unequal parental means are presented in Section S3.3 of the Supplementary Materials. The results presented in the Supplementary Materials are similar to those presented in the main text.

**Table 1:** Simulation results with weak signal strength when *G* = 10000, *H* = 150: average FDR at the nominal level 0.05 and average partial AUC (out of the maximum AUC for FPR in [0, 0.2]) for 2sLRT, Poisson MPH test, and the Point Estimation method, evaluated across two heterosis patterns.

| Metrics | Methods | Pattern A | Pattern B |
| --- | --- | --- | --- |
| FDR | 2sLRT | 0.0177 | 0.0446 |
|  | Poisson | 0.4338 | 0.9029 |
| pAUC | 2sLRT | 0.9005 | 0.8967 |
|  | Poisson | 0.8731 | 0.8741 |
|  | Estimation | 0.8227 | 0.8232 |

**Figure 2.**
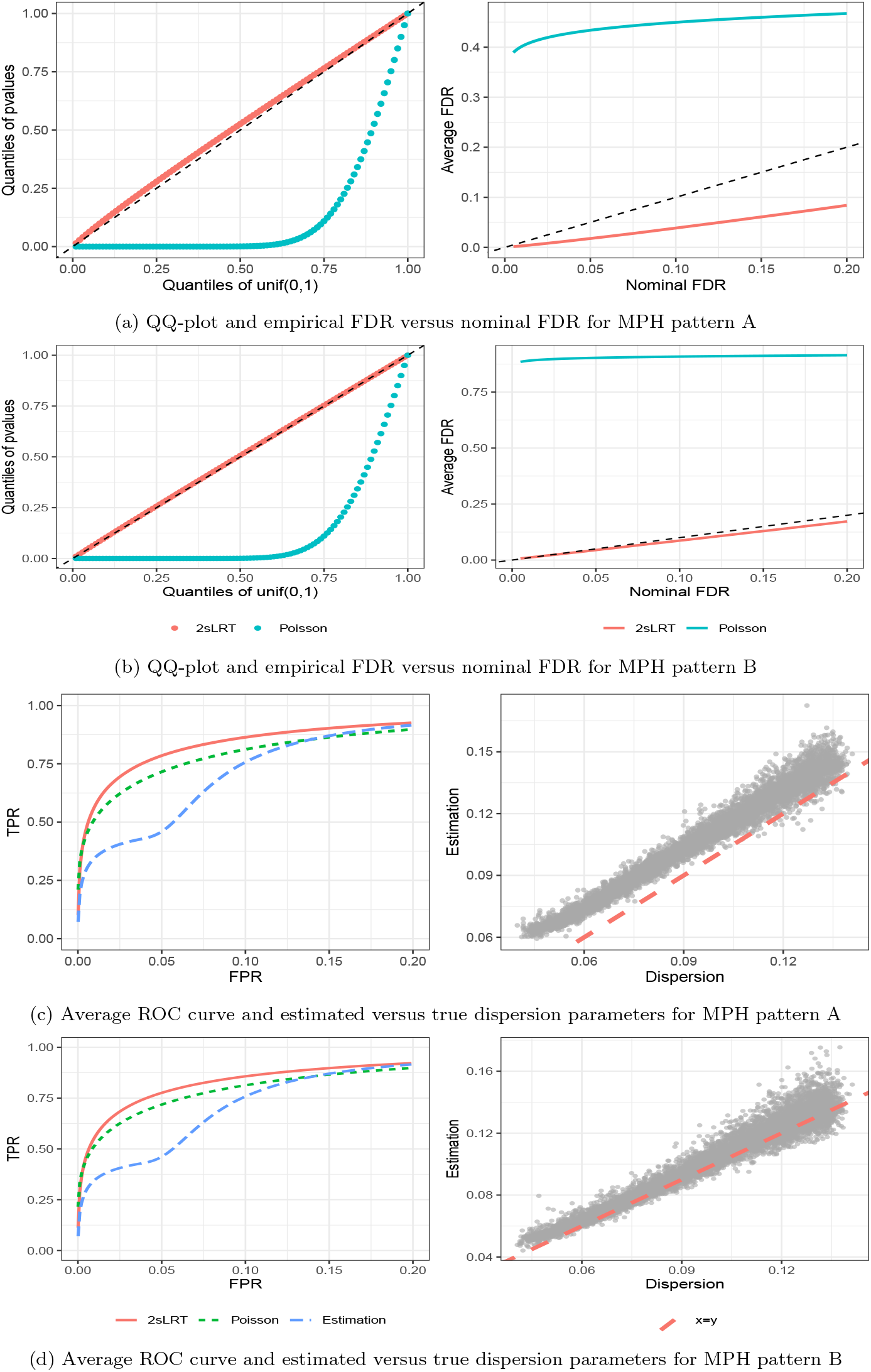
Simulation results for weak signal strength when *G* = 10000, *H* = 150 under both heterosis patterns.

We first evaluated the validity and error control of the proposed 2sLRT and the Poisson MPH test. To assess validity, we examined whether the *p*-values from non-MPH gene-family combinations followed a Uniform(0,1) distribution. As shown in Figure 2(a,b) and Supplementary Figures S1–S8, the *p*-values from 2sLRT closely followed the expected Uniform(0,1) distribution, confirming that 2sLRT produces valid tests. In contrast, the Poisson MPH test yielded *p*-values stochastically smaller than Uniform(0,1), with an excess of small values that can inflate false positives. Consistent with this, evaluation of FDR control showed that the Poisson MPH test failed to maintain FDR at the nominal level, resulting in substantial inflation of false discoveries. In comparison, 2sLRT achieved reliable FDR control under all conditions, with empirical FDR consistently below the nominal level for heterosis pattern A and close to the nominal level for pattern B (Figure 2(a,b), Table 1).

Receiver operating characteristic (ROC) curves, plotting the true positive rate (TPR) versus the false positive rate (FPR), are presented in Figure 2(c, d) and Figures S1–S8. Each curve represents averages over 50 simulated datasets, with the FPR range restricted between 0 and 0.2, where performance differences are most practically relevant. We also computed the area under the curve (AUC) and the partial AUC (pAUC), defined as the ratio of the average AUC to the maximum possible AUC within this range (Table 1). Across all settings, 2sLRT achieved the highest ROC curves and the largest pAUC values, indicating superior power at low false positive rates. Taken together with the FDR results, these findings show that 2sLRT not only ranks MPH genes more effectively but also provides more accurate FDR estimation, yielding more reliable detection of MPH genes than alternative methods.

We further evaluated the accuracy of dispersion parameter estimation using Algorithm 1. As shown in Figure 2(d), the estimated dispersion parameters closely matched the true values for heterosis pattern B, where the proportion of MPH families per gene was simulated based on empirical data. This demonstrates that the estimation procedure is robust and performs well under realistic conditions. Some overestimation was observed for pattern A, likely due to the high proportion of strong MPH signals (90% MPH families for half of the genes), which may account for the relatively conservative FDR control observed in those simulations.

The simulation results presented so far are based on *N*_0_ = 20 for the smaller dataset (*G* = 10000, *H* = 150) and *N*_0_ = 80 for the larger dataset (*G* = 30000, *H* = 600), where *N*_0_ is the threshold for the number of null families required to retain a gene in the clustering step of Algorithm 1. To evaluate the sensitivity of 2sLRT to this threshold, we examined performance under varying *N*_0_ values (Table 2), reporting the average FDR at a nominal level of 0.05 and the partial AUC for large MPH effects. Reasonable *N*_0_ values are constrained by practical limits, as overly large thresholds leave too few genes for dispersion estimation. Beyond certain points (*N*_0_ = 25 for *G* = 10000 and *N*_0_ = 90 for *G* = 30000), FDR begins to inflate and AUC declines, with the degree of degradation depending on the heterosis pattern and dataset size. In pattern A, FDR remains below nominal, whereas in pattern B, the empirical FDR exceeds the nominal level. Overall, increasing *N*_0_ initially enhances dispersion estimation by improving sample size, but excessively large values reduce the pool of genes and limit information borrowing, leading to less accurate loess fitting and more extrapolation, and compromising accuracy, especially FDR control. A well-chosen *N*_0_ balances between accuracy and stability and can be guided by inspecting the Poisson null family LRT indicator matrix.

**Table 2:** Sensitivity analysis of *N*_0_: average FDR (at the nominal level of 0.05) and pAUC (for FPR in [0,0.2]) of 2sLRT across varying numbers of gene-family combinations and two heterosis patterns under strong signal strength.

| $(G, H)$ | $N_0$ | FDR | | pAUC | |
| --- | --- | --- | --- | --- | --- |
|  |  | Pattern A | Pattern B | Pattern A | Pattern B |
| 10000, 150 | 10 | 0.0160 | 0.0371 | 0.9587 | 0.9303 |
|  | 15 | 0.0149 | 0.0362 | 0.9599 | 0.9471 |
|  | 20 | 0.0143 | 0.0282 | 0.9620 | 0.9595 |
|  | 25 | 0.0194 | 0.0393 | 0.9606 | 0.9607 |
|  | 30 | 0.0498 | 0.1022 | 0.9442 | 0.9577 |
| 30000, 600 | 60 | 0.0125 | 0.0333 | 0.9618 | 0.9503 |
|  | 70 | 0.0124 | 0.0352 | 0.9625 | 0.9581 |
|  | 80 | 0.0131 | 0.0414 | 0.9626 | 0.9601 |
|  | 90 | 0.0119 | 0.0479 | 0.9628 | 0.9614 |
|  | 100 | 0.0252 | 0.0974 | 0.9556 | 0.9544 |

Another parameter in the clustering step of Algorithm 1 is *K*, the number of clusters. We initially set *K* = 3 across all settings. While this choice produced three well-balanced clusters for the smaller dataset, it resulted in a highly imbalanced partition for the larger dataset, with most genes grouped into a single cluster. Moreover, varying *K* between 1 and 3 had little impact on overall performance. To achieve more stable dispersion estimation, we therefore set *K* = 1 for the larger dataset. In general, the choice of *K* should depend on both *N*_0_ and the similarity of null family patterns among the selected genes: more heterogeneous patterns justify a larger number of clusters, whereas more homogeneous patterns—such as those observed in our simulations—support using fewer clusters. More practical guidance on selecting *N*_0_ and *K* is discussed in Section 5.

## 4 Real Data Analysis

To study mid-parent heterosis in maize at the transcriptomic level, Xu et al. (2025) conducted a large-scale RNA-seq experiment with 802 maize varieties, including 599 hybrids derived from 203 unique parental lines. After removing genes with zero counts in any variety, 20,284 genes were retained for MPH analysis. The biological interpretations of results based on our analysis are presented in detail in Xu et al. (2025). Here, we focus on the statistical analysis and the methodological aspect of the study.

We applied the 2sLRT to detect MPH genes with the following settings in our algorithms: the FDR threshold for identifying “null” families was set to 0.1, *N*_0_ to 50, the number of clusters *K* to 10, and the loess span to 0.1. We first examined the gene-wise dispersion parameter estimates. As shown in Supplementary Figure S11, these estimates exhibited a clear inverse relationship with the median mid-parent expression levels, consistent the trends observed in other RNA-seq datasets. This pattern was well captured by a loess fit, and the resulting estimates were constrained within a reasonable range, supporting the validity of our estimation procedure. Finally, *p*-values from both the 2sLRT and the Poisson MPH test were subject to FDR control at a nominal level of 0.05 as described in Algorithm 2 of Supplementary Materials.

Consistent with the findings from our simulations, the proposed 2sLRT exhibits markedly higher specificity in identifying true MPH gene–hybrid combinations than the Poisson MPH test, thereby substantially reducing false positives. At a significance level of 0.05 (based on adjusted *p*-values), 2sLRT identifies 6.81% of gene–family combinations as exhibiting MPH, compared with 63.43% identified by the Poisson MPH test. The excessive number of significant results from the Poisson test mirrors its inflated false positive rate observed in simulations and contrasts with prior studies indicating that mid-parent heterosis is limited to a small subset of genes in maize and other organisms.

To further assess specificity, we examined a set of nine housekeeping genes (Lin et al., 2014), which are characterized by stable expression across varieties and are not expected to exhibit MPH. As shown in Table 3, 2sLRT reports the proportion of MPH hybrids for these genes to be close to zero, indicating accurate classification. In contrast, the Poisson MPH test falsely identifies 50% to 78% of hybrids as exhibiting MPH for these genes. These results reinforce the superior error control and reliability of 2sLRT in real data analysis.

**Table 3:** Proportion of MPH hybrids for nine housekeeping genes, based on 2sLRT or Poisson MPH test adjusted *p*-values at a significance level of 0.05.

| Housekeeping genes | Proportion of MPH hybrids |  |
| --- | --- | --- |
|  | 2sLRT | Poisson MPH test |
| Zm00001d045417 | 0.02 | 0.511 |
| Zm00001d010476 | 0.018 | 0.641 |
| Zm00001d015962 | 0.007 | 0.596 |
| Zm00001d039138 | 0.03 | 0.778 |
| Zm00001d051879 | 0.023 | 0.591 |
| Zm00001d002944 | 0.003 | 0.533 |
| Zm00001d044172 | 0.013 | 0.751 |
| Zm00001d006438 | 0.077 | 0.758 |
| Zm00001d036201 | 0.002 | 0.474 |

We next examined the patterns among detected MPH gene-family combinations by 2sLRT. Our analysis reveals that the gene-wise MPH pattern is much more pronounced than the hybrid-wise pattern. The left panel of Figure 3 shows the histogram of the proportions of MPH hybrids for the 20284 genes. The variation in proportions of MPH hybrids across different genes is substantial, spanning from 0 to 0.9, and the distribution is concentrated towards zero and has a heavy right tail. This suggests that most genes rarely show the MPH pattern across hybrids, while a subset exhibits MPH broadly across many hybrids. In contrast, the right panel shows the histogram of MPH gene proportions for the 599 hybrids. Here, the range of MPH gene proportions across hybrids is considerably narrower than the range of MPH hybrid proportions across genes, and the distribution is smoother, indicating less variability in the number of MPH genes per hybrid. This pattern implies that heterosis may be driven by a specific set of genes dominant across genetic backgrounds, rather than being a highly hybrid-specific phenomenon. Additional visualizations supporting this data analysis are provided in Section S4 of the Supplementary Materials.

**Figure 3.**
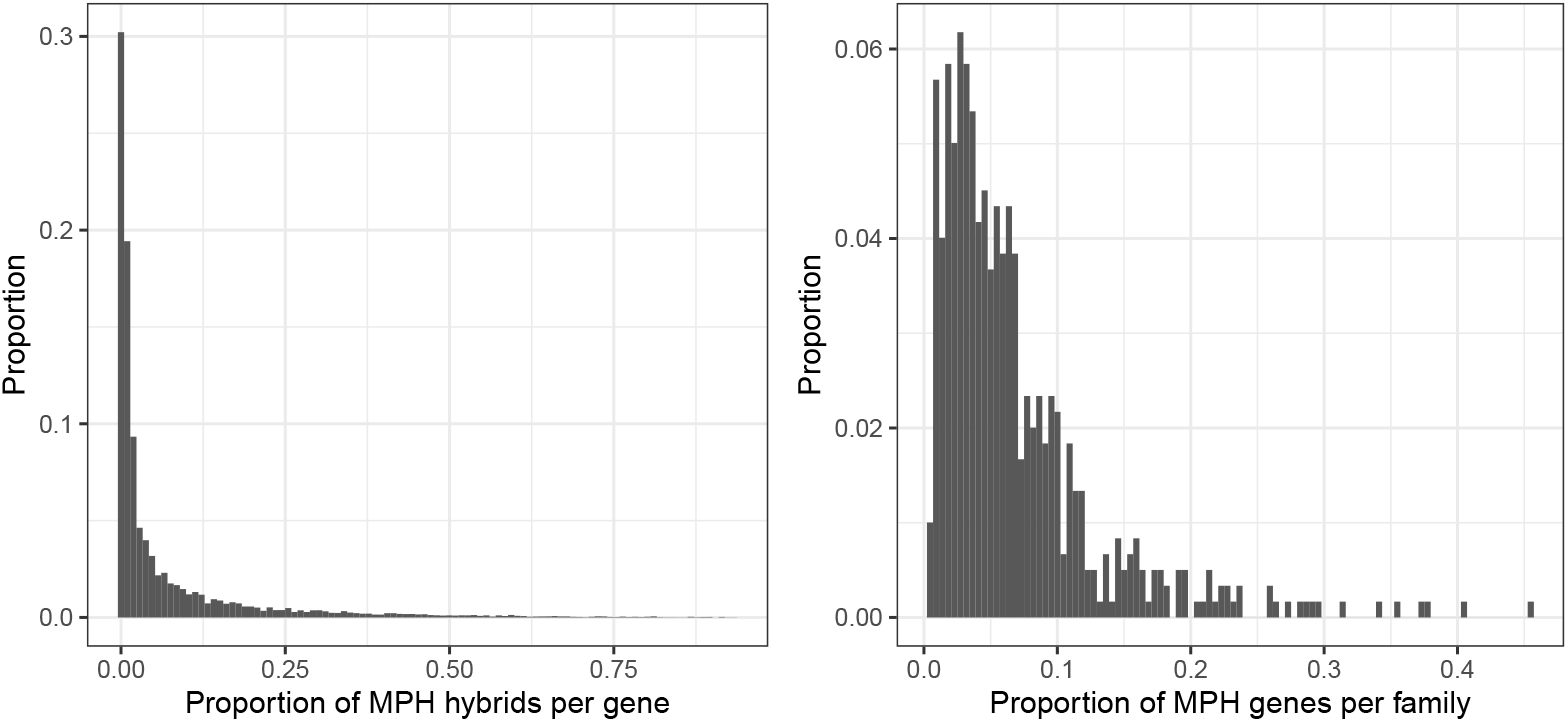
Histogram of gene-wise MPH hybrids proportions (left) and family-wise MPH gene proportions (right).

## 5 Discussion

In this study, we developed a novel two-stage likelihood ratio test (2sLRT) to detect MPH genes in large-scale unreplicated RNA-seq experiments. The proposed method specifically addresses the key challenge of estimating dispersion parameters in the negative binomial model under limited replication, thereby enabling more reliable and accurate assessment of MPH patterns than existing approaches.

Accurate estimation of dispersion parameters is central to negative binomial–based inference, including the detection of MPH genes. In our 2sLRT framework, several algorithmic parameters influence estimation accuracy, among which *N*_0_, the minimum number of null families required for a gene to be included in the clustering step, is important. Increasing *N*_0_ enlarges the null family set and provides more information for dispersion estimation; however, our sensitivity analysis indicates that setting *N*_0_ too high reduces the number of eligible genes, limiting information borrowing and increasing the risk of over-extrapolation. In practice, *N*_0_ should be selected by evaluating multiple candidate values and choosing one that achieves strong within-cluster agreement among null families while retaining genes that span a broad range of expression levels. This can be assessed by examining the sub-indicator matrices from the Poisson null-family LRT and inspecting plots of median mid-parent expression versus both estimated and predicted dispersion values.

The number of clusters *K* is another algorithmic parameter and should be selected in conjunction with *N*_0_. A larger *K* is appropriate when genes exhibit heterogeneous null-family patterns, whereas fewer clusters, or even a single cluster, are sufficient when genes are largely homogeneous. In practice, users may begin with a relatively large *K*, assess the resulting cluster sizes and homogeneity, and then merge clusters as needed to ensure each cluster contains enough genes and maintains sufficient homogeneity for reliable dispersion estimation.

Using the above strategies to select *N*_0_ and *K*, our simulation studies demonstrate that the 2sLRT effectively controls the FDR while maintaining strong statistical power. Across all scenarios, the area under the ROC curve for 2sLRT consistently exceeds that of both the Poisson MPH test and the naïve estimation approach. Furthermore, application to a maize dataset with 599 hybrids showed that 2sLRT markedly reduces false positives, confirming its reliability and value for large-scale transcriptomic studies.

## Data Availability

The raw sequence data reported in this paper have been deposited in the Genome Sequence Archive in the National Genomics Data Center, China National Center for Bioinformation, Beijing Institute of Genomics, Chinese Academy of Sciences (GSA: CRA023776), which are publicly accessible at https://ngdc.cncb.ac.cn/gsa. An R package for implementing 2sLRT can be accessed from the GitHub repository https://github.com/yunhuiqistat/TwoStageLRT.

