## Supplementary materials for "Detection of Mid-parent Heterosis Genes in Large-Scale Unreplicated RNA-Seq Experiments"

Yunhui Qi*

Department of Data Science, Dana-Farber Cancer Institute

and

Peng Liu †

Department of Statistics, Iowa State University

December 11, 2025

*Contributing author:

**S4 Additional Figures for Example Data Analysis** **10**

### List of Tables

S1 Simulation results with weak signal strength for $G = 30000$ and $H = 600$ . . 8

S2 Simulation results with strong signal strength . . . . . . . . . . . . . . . . . 8

S3 Simulation results when parental means are unequal with strong signal strength 16

S4 Simulation results when parental means are unequal with weak signal strength 16

S11 Dispersion estimation versus median mid-parent mean in example data analysis 18

### S1 Algorithms

**Algorithm 1** Gene-wise dispersion parameter estimation

**Require:** RNA-seq count data for $G$ genes and $H$ families. Minimum number of null families for each gene to estimate dispersion parameter $N_0$, threshold of FDR for construction of indicator matrix $\alpha_0$, number of clusters $K$, normalization factors $C_{hi}$ for $h = 1, \ldots, H$ and $i = 1, 2, 3$, and span for loess curve $s$.

**1. Poisson null-family likelihood ratio test:** For each gene and family combination, perform Poisson LRT for differential expression analysis conditional on the normalization factors. Adjust the resulting $p$-values for FDR control using Benjamini-Hochberg method. Get the FDR matrix $\mathbf{Q} = (q_{gh})_{g=1,\ldots,G}^{h=1,\ldots,H}$ and an indicator matrix $\mathbf{I} = (i_{gh}) = 1(q_{gh} > \alpha_0)$.

**2. Clustering of genes:** Conduct hierarchical clustering of genes with more than $N_0$ null families using complete linkage, with the Jaccard distance computed from the indicator matrix $\mathbf{I}$, and partition the genes into $K$ clusters.

**3. Estimation of dispersion parameters:**

**for** each cluster of genes **do**

Find the gene with the smallest number of null families. Denote the set of indices of null families for this gene as $\mathbf{M}$.

Estimate the gene-wise dispersion parameters using edgeR on families in $\mathbf{M}$ and all genes in this cluster.

**end for**

**4. Prediction and extrapolation:**

Fit a loess curve with span $s$ using median mid-parent mean as the predictor and the estimated dispersion parameters as the response.

For genes with fewer than $N_0$ null families, predict their dispersion parameters using the fitted loess curve.

For genes with fewer than $N_0$ null families and with median mid-parent mean outside the fitted loess curve, assign the dispersion parameter from the gene with the nearest median mid-parent mean.

**return** Dispersion parameter value for each gene of $G$ genes.

**Algorithm 2** Detection of MPH genes

**Require:** RNA-seq count data for $G$ genes and $H$ families, where each family contains one hybrid variety and two parental varieties with a single biological replicate for each variety. Minimum number of null families for each gene to estimate dispersion parameter $N_0$, threshold of FDR for construction of indicator matrix $\alpha_0$, number of clusters $K$, and span for loess curve $s$.

**1. Normalization:** Apply methods such as DESeq or TMM to all samples to get the normalization factors.

**2. Estimation of dispersion parameters:** Use Algorithm 1 to get gene-wise dispersion estimates for $G$ genes given $N_0$, $K$, $\alpha_0$, $s$, and normalization factors. Denote the estimated dispersion parameters as $\{d_g, g = 1, \ldots, G\}$.

**3. Negative binomial likelihood ratio test conditional on estimated dispersion parameters:** For each gene and family combination, perform the NB LRT with the estimated dispersion parameters and normalization factors, and get the $p$-value matrix $\mathbf{P}$.

**4. False discovery rate control:** Apply Benjamini-Hochberg method to each column of $\mathbf{P}$ to get the FDR matrix $\mathbf{Q}$.

**return** FDR matrix for $G$ genes and $H$ families for mid-parent heterosis.

### S2 Derivation of Poisson LRT for Mid-parent Heterosis

To detect MPH genes, we need to test the hypotheses below.

$$H_{gh0} : \mu_{gh3} = (\mu_{gh1} + \mu_{gh2})/2 \text{ versus } H_{gh1} : \mu_{gh3} \neq (\mu_{gh1} + \mu_{gh2})/2. \tag{S1}$$

The Poisson LRT for testing $H_{gh0}$ is derived as follows. Following the notations in Section 2, suppose $y_{ghi} \sim \text{Poisson}(C_{hi}\mu_{ghi})$, the MLE of $\mu_{ghi}$ are $\hat{\mu}_{ghi} = y_{ghi}/C_{hi}$ under $H_{gh1}$. Under $H_{gh0}$, MLE of $\mu_{ghi}$ should satisfy

$$\begin{aligned} \frac{y_{gh1}}{\hat{\mu}^0_{gh1}} + \frac{y_{gh3}}{\hat{\mu}^0_{gh1} + \hat{\mu}^0_{gh2}} - C_{h1} - \frac{C_{h3}}{2} &= 0 \\ \frac{y_{gh2}}{\hat{\mu}^0_{gh2}} + \frac{y_{gh3}}{\hat{\mu}^0_{gh1} + \hat{\mu}^0_{gh2}} - C_{h2} - \frac{C_{h3}}{2} &= 0 \\ \hat{\mu}^0_{gh1} + \hat{\mu}^0_{gh2} &= 2\hat{\mu}^0_{gh3}. \end{aligned} \tag{S2}$$

Solving the system of equations (S2) reduces to finding the positive root of a quadratic equation, which yields a unique solution. This allows us to compute the MLE under $H_{gh0}$ and perform the Poisson LRT for MPH.

As a special case, when $C_{hi} = 1, i = 1, 2, 3$, under $H_{gh0}$, MLE of $\mu_{ghi}, i = 1, 2, 3$ are:

$$\hat{\mu}^0_{gh1} = \frac{2y_{gh1}(y_{gh1} + y_{gh2} + y_{gh3})}{3(y_{gh1} + y_{gh2})}, \hat{\mu}^0_{gh2} = \frac{2y_{gh2}(y_{gh1} + y_{gh2} + y_{gh3})}{3(y_{gh1} + y_{gh2})}, \hat{\mu}^0_{gh3} = \frac{1}{3}(y_{gh1}+y_{gh2}+y_{gh3}).$$

Globally, the MLEs are

$$\hat{\mu}^a_{gh1} = y_{gh1}, \hat{\mu}^a_{gh2} = y_{gh2}, \hat{\mu}^a_{gh3} = y_{gh3}.$$

Then the test statistic for Poisson LRT is

$$R^p_{gh} = -2\left[(y_{gh1} + y_{gh2}) \log \frac{2(y_{gh1} + y_{gh2} + y_{gh3})}{3(y_{gh1} + y_{gh2})} + y_{gh3} \log \frac{y_{gh1} + y_{gh2} + y_{gh3}}{3y_{gh3}}\right].$$

The null hypothesis will be rejected if $R^p_{gh} > \chi^2_1(1 - \alpha)$.

### S3 Additional Simulation Results

#### S3.1 Simulation settings

We provide a visual aid for the simulation design described in Section 3 of the main text. Figure S1 illustrates the hierarchical workflow used to simulate RNA-seq data in panel (a) and details the parameter settings for the two distinct MPH patterns investigated in panel (b).

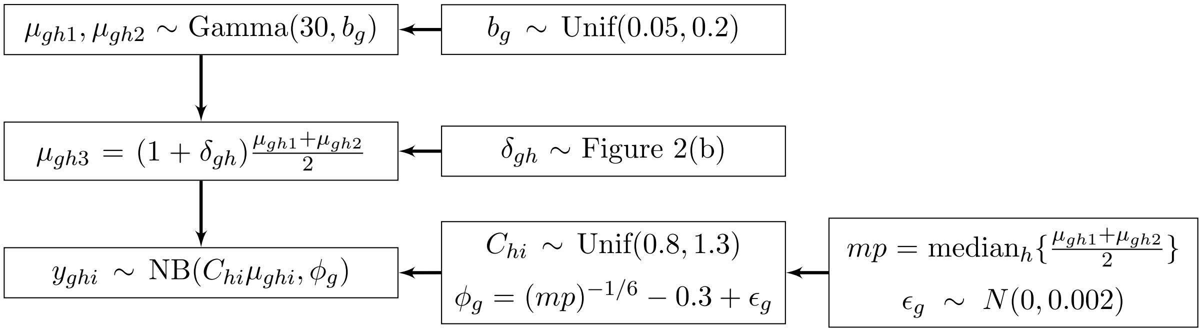

**(a)**

| Pattern A | | Pattern B | |
|---|---|---|---|
| Proportion of genes | MPH effect | Proportion of genes | MPH effect |
| 1/4 | $\delta_{gh} > 0$ in 5% families | 1/2 | $\forall g$, $p_g \sim$ empirical distribution, $\delta_{gh} < 0$ in $p_g$% families |
| 1/4 | $\delta_{gh} < 0$ in 5% families | | |
| 1/4 | $\delta_{gh} > 0$ in 90% families | 1/2 | $\forall g$, $p_g \sim$ empirical distribution, $\delta_{gh} > 0$ in $p_g$% families |
| 1/4 | $\delta_{gh} < 0$ in 90% families | | |

**(b)**

Figure S1: Simulation overview. (a) Workflow for simulating RNA-seq data, illustrating the hierarchical structure and key parameter settings. (b) Simulation settings defining two patterns of mid-parent heterosis (MPH): Pattern A with fixed proportions of MPH families, and Pattern B with empirical, gene-specific proportions of MPH families. $\delta_{gh}$ = 2 or 3 when $\delta_{gh} > 0$, and $\delta_{gh}$ = -2/3 or -3/4 when $\delta_{gh} < 0$.

#### S3.2 Equal parental mean

We first present simulation results for the case where parental means are simulated to be equal. Table S1 summarizes the average FDR and pAUC, while Figure S2 and Figure S3 show the QQ plots of $p$-values for null genes, empirical versus nominal FDR, average ROC curves, and the relationship between estimated and true dispersion parameters under weak signal strength ($G = 30000, H = 600$) for heterosis patterns A and B, respectively.

Simulation results under strong signal strength are summarized in Table S2. Corresponding figures for heterosis patterns A and B are shown in Figure S4 and Figure S5 for the smaller dataset ($G = 10000, H = 150$), and in Figure S6 and Figure S7 for the larger dataset ($G = 30000, H = 600$).

| | | $G = 30000, H = 600$ | |
|---|---|---|---|
| Metrics | Methods | Pattern A | Pattern B |
| FDR | 2sLRT | 0.0169 | 0.0464 |
| | Poisson | 0.4341 | 0.9050 |
| pAUC | 2sLRT | 0.9007 | 0.8983 |
| | Poisson | 0.8731 | 0.8736 |
| | Estimation | 0.8213 | 0.8228 |

Table S1: Simulation results with weak signal strength for $G = 30000$ and $H = 600$: average FDR at nominal FDR level 0.05 and average partial AUC (FPR in [0, 0.2]) for 2sLRT, Poisson MPH test, and the Point Estimation method, evaluated across different gene and family combinations and two heterosis patterns.

| | | $G = 10000, H = 150$ | | $G = 30000, H = 600$ | |
|---|---|---|---|---|---|
| Metrics | Methods | Pattern A | Pattern B | Pattern A | Pattern B |
| FDR | 2sLRT | 0.0142 | 0.0358 | 0.0131 | 0.0414 |
| | Poisson | 0.4320 | 0.9029 | 0.4316 | 0.9053 |
| pAUC | 2sLRT | 0.9618 | 0.9591 | 0.9626 | 0.9601 |
| | Poisson | 0.9304 | 0.9309 | 0.9308 | 0.9308 |
| | Estimation | 0.8819 | 0.8804 | 0.8827 | 0.8814 |

Table S2: Simulation results with strong signal strength: average FDR at nominal FDR level 0.05 and average partial AUC (FPR in [0, 0.2]) for 2sLRT, Poisson MPH test, and the Point Estimation method, evaluated across different gene and family combinations and two heterosis patterns.

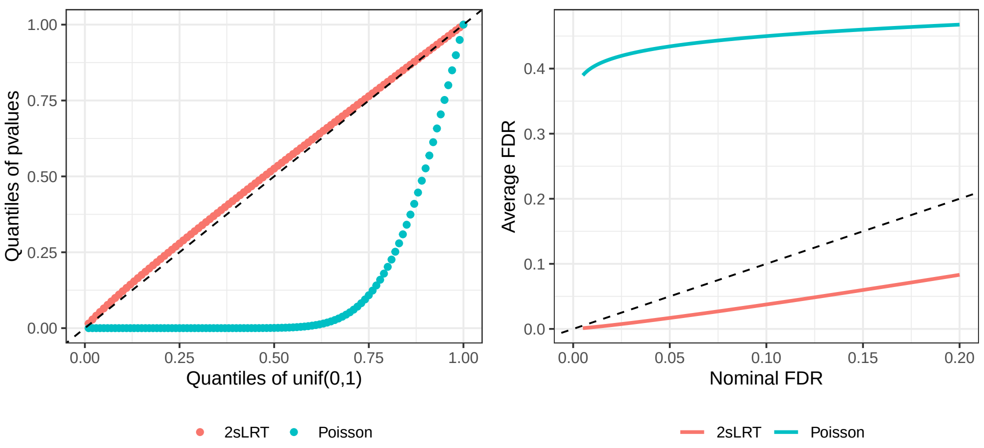

(a) QQ-plot and empirical FDR versus nominal FDR

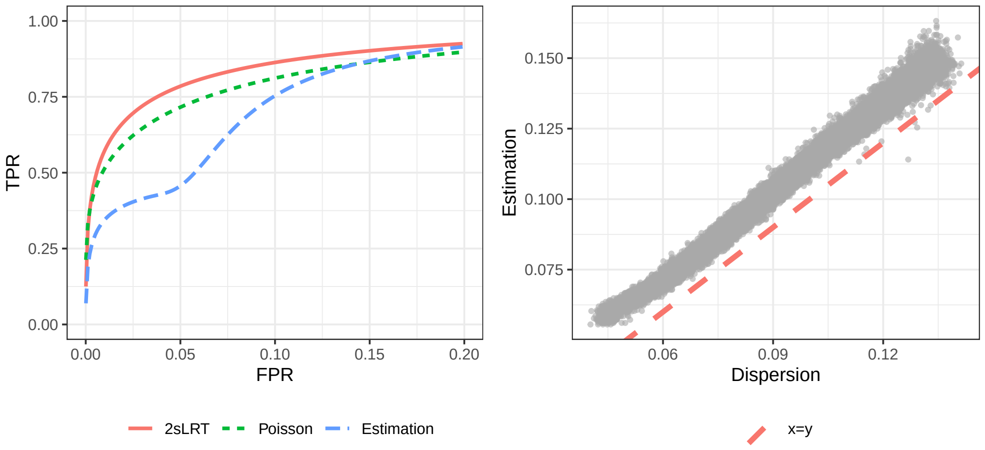

(b) Average ROC curve and estimated versus true dispersion parameters

Figure S2: Simulation results for weak signal strength when $G = 30000, H = 600$ under heterosis pattern A.

#### S3.3 Unequal parental mean

We also present simulation results where parental means are simulated independently, with unequal means for the second half of the genes. In these simulations (Figure 2), $\mu_{gh1}$ is generated as described in Section 3. For $\mu_{gh2}$, we set $\mu_{gh2} = \mu_{gh1}$ for $g = 1, \ldots, G/2$ to retain some null families, and simulate $\mu_{gh2}$ for $g = G/2 + 1, \ldots, G$ from Gamma$(30, b_g)$ to ensure unequal parental means for the remaining genes.

Results for strong and weak signals are summarized in Tables S3 and S4, respectively. The corresponding QQ plots, empirical versus nominal FDR, ROC curves, and scatter plots

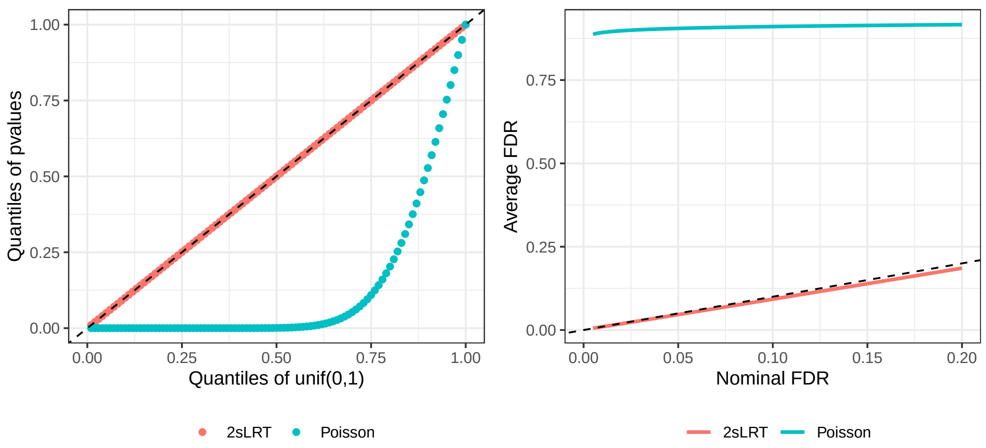

(a) QQ-plot and empirical FDR versus nominal FDR

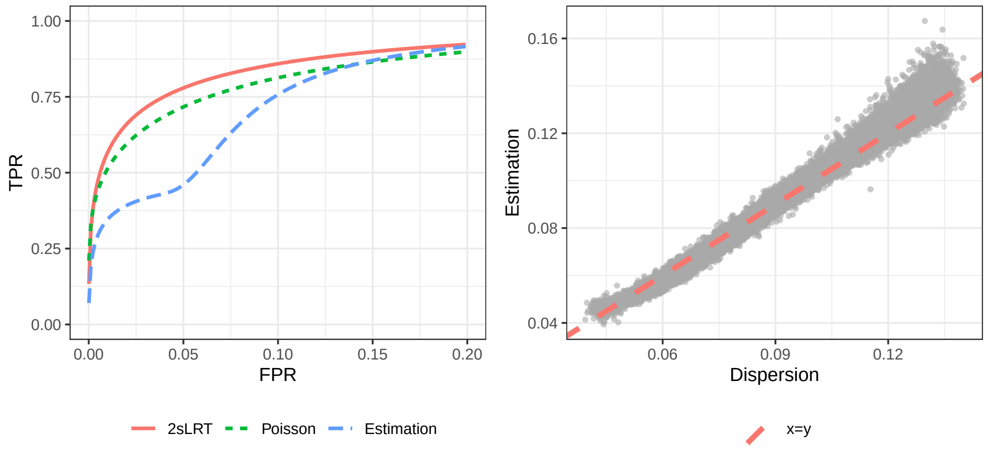

(b) Average ROC curve and estimated versus true dispersion parameters

Figure S3: Simulation results for weak signal strength when $G = 30000, H = 600$ under heterosis pattern B.

of estimated versus true dispersion parameters are shown in Figures S8 (strong signal) and S9 (weak signal). The results are consistent with those in Section 3, confirming the superior power and FDR control of the proposed method over other methods.

### S4 Additional Figures for Example Data Analysis

Figure S10 shows the $p$-value distributions from the Poisson MPH test and the proposed 2sLRT. The results are consistent with the results that we present in the main text, that 2sLRT identified much fewer MPH gene–hybrid combinations than the Poisson MPH test,

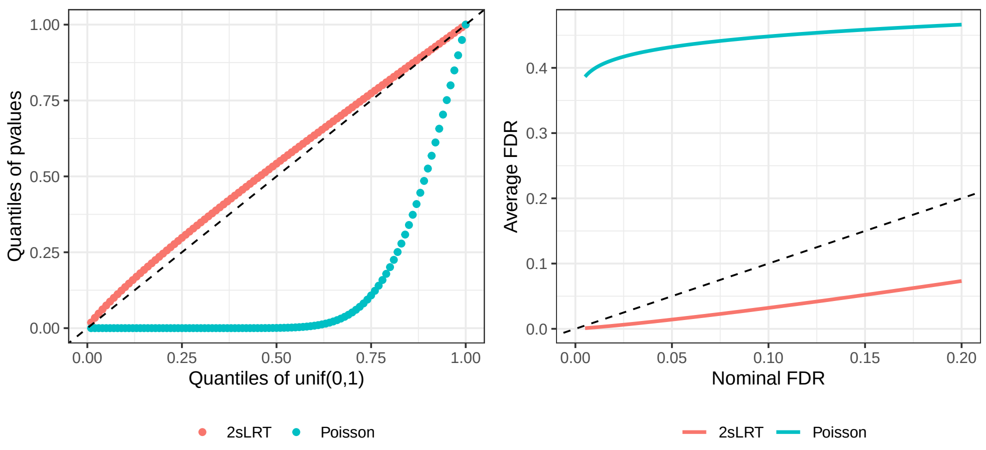

(a) QQ-plot and empirical FDR versus nominal FDR

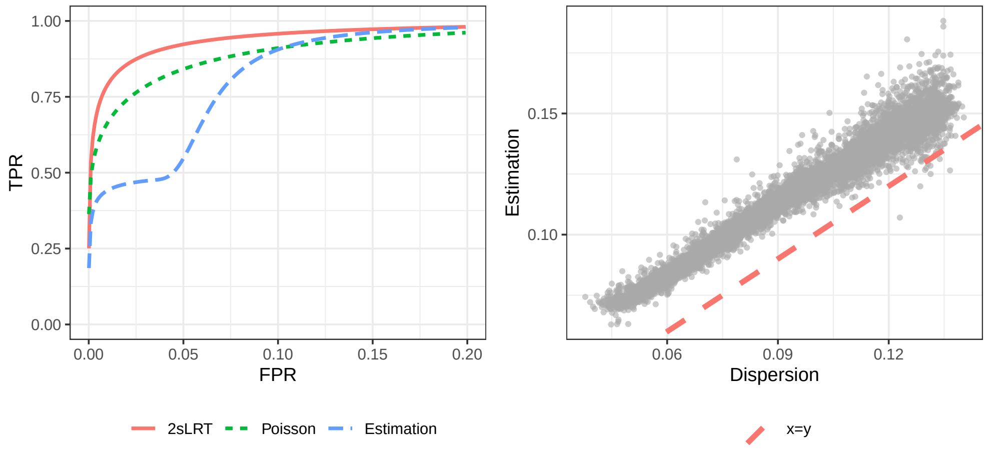

(b) Average ROC curve and estimated versus true dispersion parameters

Figure S4: Simulation results for strong signal strength when $G = 10000, H = 150$ under heterosis pattern A.

which tends to produce many false positives.

Figure S11 displays the estimated dispersion parameters from 2sLRT plotted against the median mid-parent mean for 20,284 genes across 599 families. Each black dot represents an estimated dispersion value, which ranges from 0 to 0.9. As the median mid-parent mean increases, both the dispersion estimates and their variability decrease, similar to the patterns observed in other datasets. The red curve shows the fitted LOESS trend, and the green points denote dispersion values predicted from this curve. Although the LOESS fit does not capture the variability at the lower range, it provides an overall satisfactory representation of the trend.

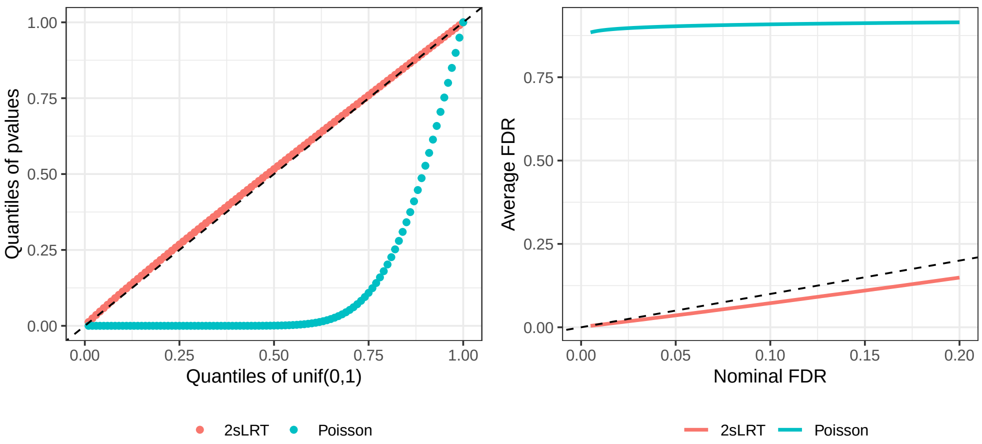

(a) QQ-plot and empirical FDR versus nominal FDR

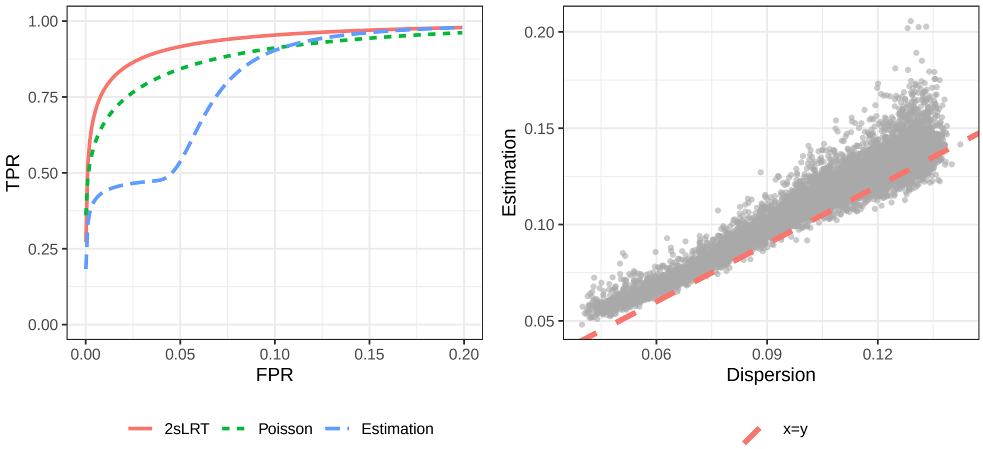

(b) Average ROC curve and estimated versus true dispersion parameters

Figure S5: Simulation results for strong signal strength when $G = 10000, H = 150$ under heterosis pattern B.

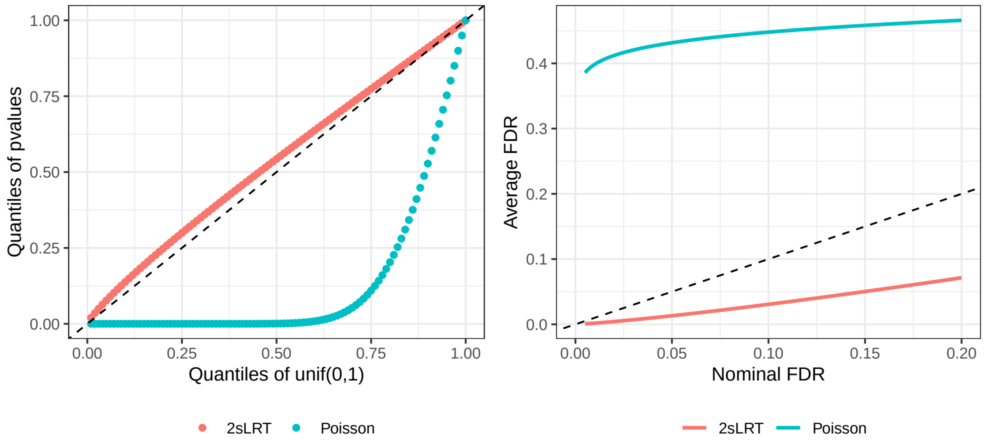

(a) QQ-plot and empirical FDR versus nominal FDR

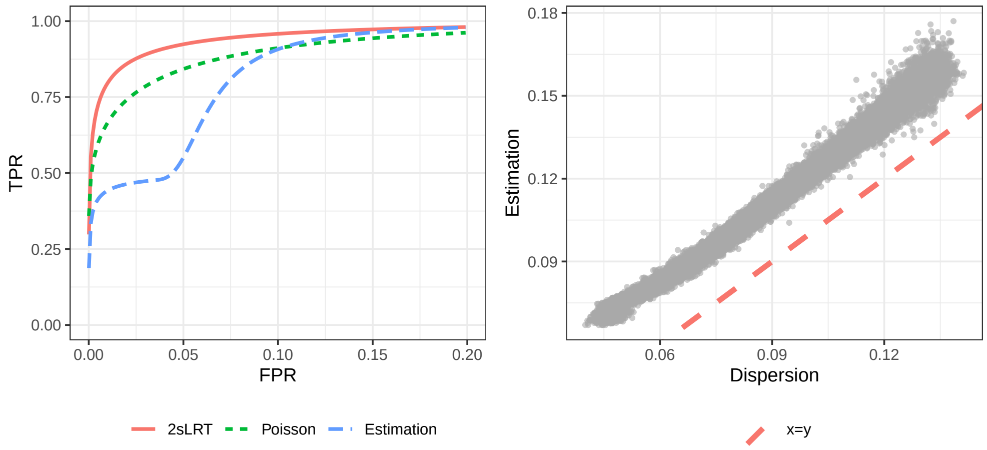

(b) Average ROC curve and estimated versus true dispersion parameters

Figure S6: Simulation results for strong signal strength when $G = 30000, H = 600$ under heterosis pattern A.

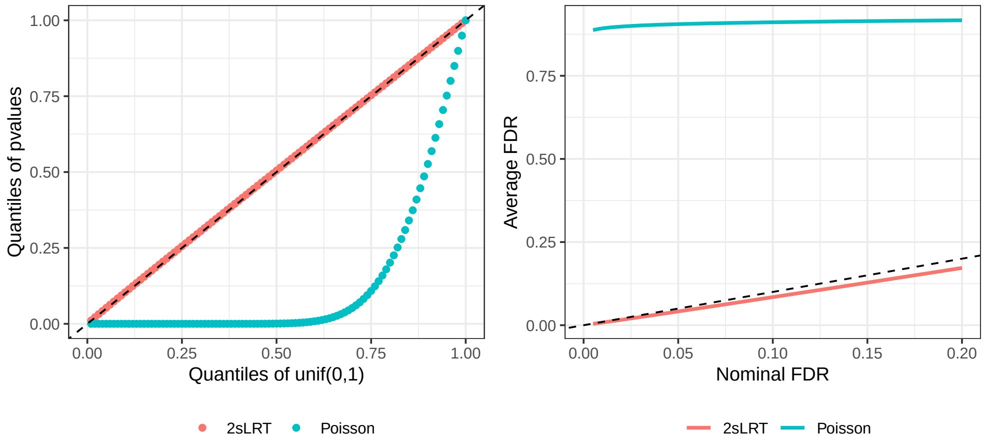

(a) QQ-plot and empirical FDR versus nominal FDR

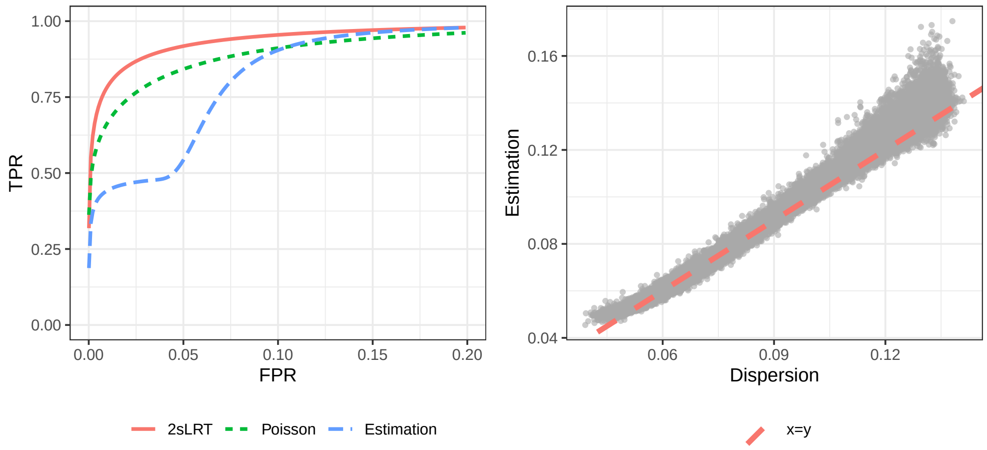

(b) Average ROC curve and estimated versus true dispersion parameters

Figure S7: Simulation results for strong signal strength when $G = 30000, H = 600$ under heterosis pattern B.

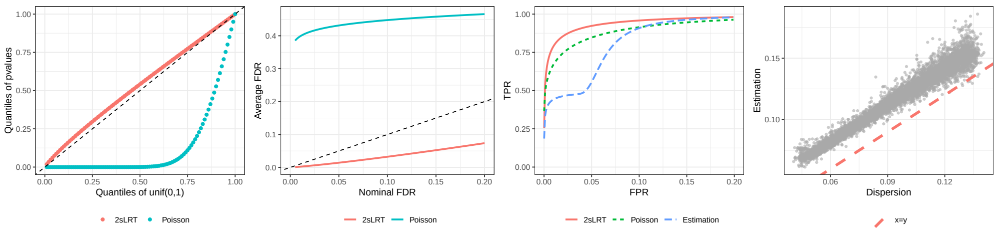

(a) $G = 10000, H = 150$ with heterosis pattern A

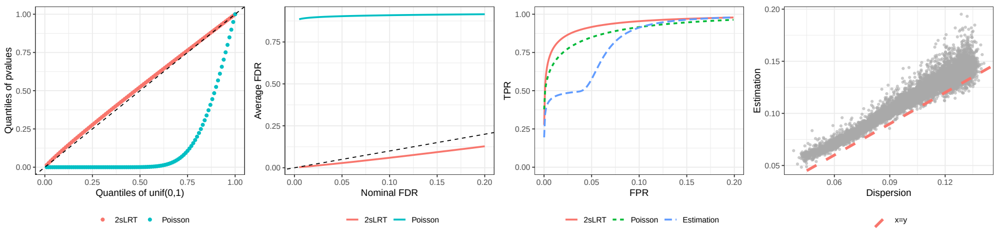

(b) $G = 10000, H = 150$ with heterosis pattern B

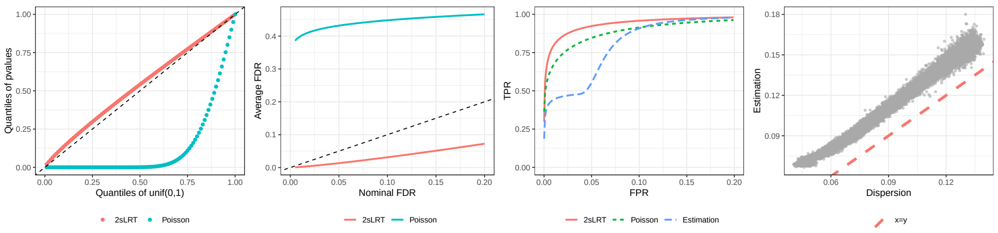

(c) $G = 30000, H = 600$ with heterosis pattern A

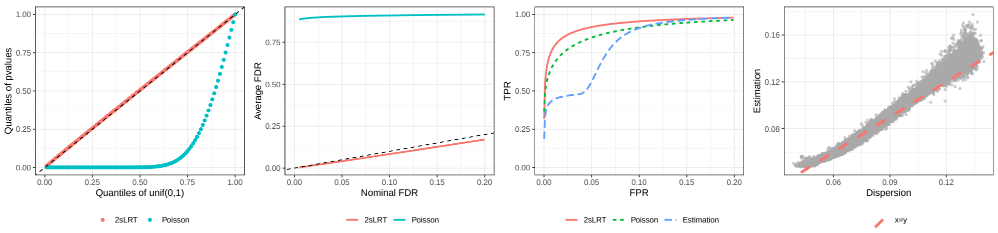

(d) $G = 30000, H = 600$ with heterosis pattern B

Figure S8: Simulation results when parental means are unequal with strong signal strength: QQ-plot, empirical FDR versus nominal FDR, average ROC curve, and scatter plot between estimated and true dispersion parameters, evaluated under different numbers of gene and family combinations and two heterosis patterns.

| **Strong signal** | | $G = 10000, H = 150$ | | $G = 30000, H = 600$ | |
|---|---|---|---|---|---|
| Metrics | Methods | Pattern A | Pattern B | Pattern A | Pattern B |
| FDR | 2sLRT | 0.0143 | 0.0282 | 0.0135 | 0.0411 |
| | Poisson | 0.4314 | 0.9037 | 0.4318 | 0.9033 |
| pAUC | 2sLRT | 0.9620 | 0.9595 | 0.9624 | 0.9601 |
| | Poisson | 0.9326 | 0.9337 | 0.9323 | 0.9334 |
| | Estimation | 0.8833 | 0.8875 | 0.8831 | 0.8844 |

Table S3: Simulation results when parental means are unequal with strong signal strength: average FDR at nominal FDR level 0.05 and average pAUC (FPR in [0,0.2]) for 2sLRT, Poisson MPH test, and the Point Estimation method under different numbers of gene and family combinations, two heterosis patterns.

| **Weak signal** | | $G = 10000, H = 150$ | | $G = 30000, H = 600$ | |
|---|---|---|---|---|---|
| Metrics | Methods | Pattern A | Pattern B | Pattern A | Pattern B |
| FDR | 2sLRT | 0.0176 | 0.0349 | 0.0166 | 0.0448 |
| | Poisson | 0.4343 | 0.8996 | 0.4346 | 0.9045 |
| pAUC | 2sLRT | 0.8998 | 0.8979 | 0.9011 | 0.8973 |
| | Poisson | 0.8754 | 0.8761 | 0.8752 | 0.8747 |
| | Estimation | 0.8223 | 0.8265 | 0.8232 | 0.8262 |

Table S4: Simulation results when parental means are unequal with weak signal strength: average FDR at nominal FDR level 0.05 and average pAUC (FPR in [0,0.2]) for 2sLRT, Poisson MPH test, and the Point Estimation method under different numbers of gene and family combinations, two heterosis patterns.

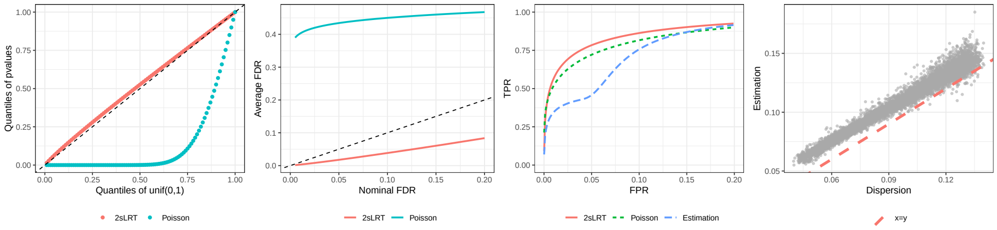

(a) $G = 10000, H = 150$ with heterosis pattern A

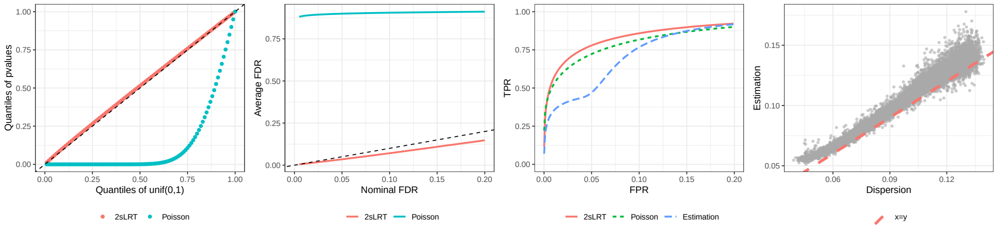

(b) $G = 10000, H = 150$ with heterosis pattern B

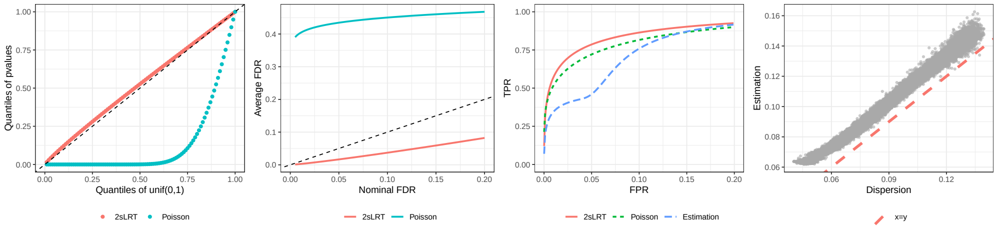

(c) $G = 30000, H = 600$ with heterosis pattern A

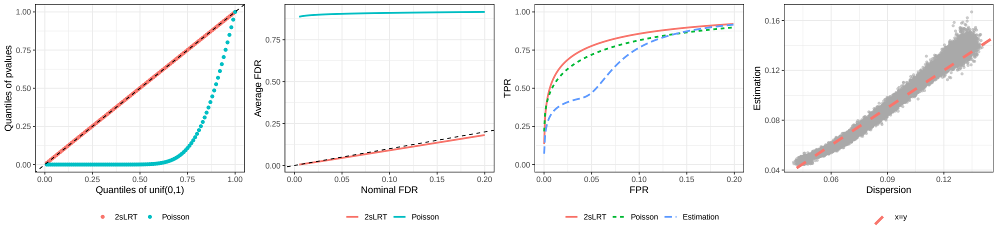

(d) $G = 30000, H = 600$ with heterosis pattern B

Figure S9: Simulation results when parental means are unequal with weak signal strength: QQ-plot, empirical FDR versus nominal FDR, average ROC curve, and scatter plot between estimated and true dispersion parameters, evaluated under different numbers of gene and family combinations and two heterosis patterns

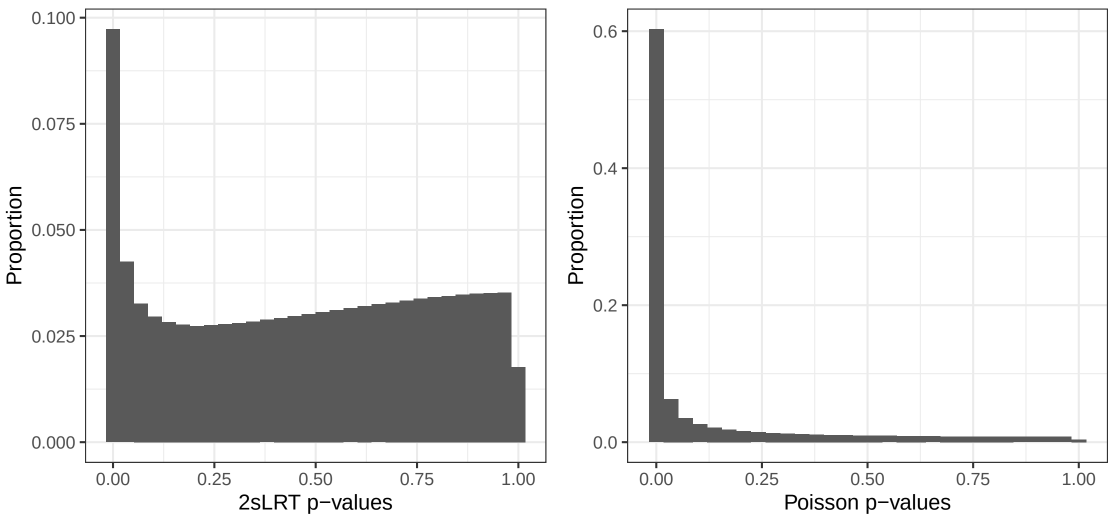

Figure S10: Histogram of $p$-values from 2sLRT and Poisson MPH test in example data analysis.

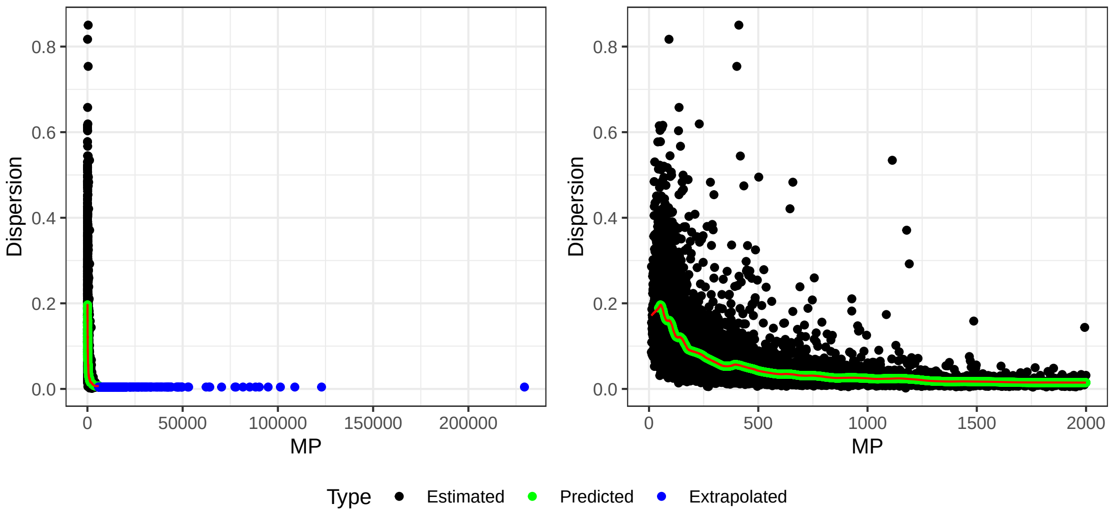

Figure S11: Relationship between dispersion estimation from 2sLRT and median of mid-parent mean estimation across 599 families for 20284 genes. The right panel is a zoom-in version of the left panel.
